# A fast TMT-based proteomic workflow reveals neural enrichment in neurospheres of hiPSC-derived neural stem cells

**DOI:** 10.1101/2025.06.09.658417

**Authors:** Pedro de Lima Muniz, Leticia Rocha Quintino Souza, Michele Rodrigues Martins, Wassali Valadares, Fábio César Sousa Nogueira, Stevens Rehen, Guillaume Nugue, Magno Junqueira

**Author notes:** Corresponding author: Magno Junqueira – Laboratório de Espectrometria de Massa Translacional e Neuroproteômica (LEMTEN), Cidade Universitária, Centro de Tecnologia, block A, 5th floor, room 543 - Rio de Janeiro 21941-909, Brazil.

## Abstract

Three-dimensional (3D) neural spheroids, or neurospheres, generated from human induced pluripotent stem cell (hiPSC)-derived neural stem cells (NSCs) more accurately recapitulate the microenvironmental cues of neural tissue compared to traditional two-dimensional (2D) monolayers. However, comparative omics-based characterizations of these models remain limited. Here, we present a streamlined and scalable TMT-based quantitative proteomics workflow to contrast the proteomic landscapes of hiPSC-derived NSCs cultured in 2D monolayers versus 3D neurospheres. A total of 1,576 proteins were identified in an unfractionated LC-MS/MS of 68 minutes, with 542 showing significant differential abundance between groups. Neurospheres exhibited enrichment in neural-related pathways, such as synaptic signaling, neurotrophin signaling, cytoskeletal organization and vesicle trafficking, while monolayers enriched for multipotency features, such as general metabolic activity. Cell-type enrichment analyses confirmed increased neuronal identity in neurospheres, including elevated levels of markers associated with neuronal maturation. Our results demonstrate that 3D culture of NSCs induces a proteomic shift toward a more mature neural phenotype. This rapid, multiplexed proteomic approach enables high-content molecular profiling suitable for drug screening and personalized medicine applications.

**Significance:** The present work contributes to the molecular and biological understanding of iPSC-derived neurospheres by exploring the proteome of this model. Neurospheres are a culture model of great potential and applicability in neurobiology research and personalized medicine, which still lacks a robust omic characterization. By studying neurospheres, we also work with a 3D culture model generated *in vitro*, avoiding the use of primary neural cells culture and animal models. Our fast method is relevant to single-cell proteomics, personalized medicine and screening assays.

## Introduction

The study of the human nervous system at a molecular or histological level is surrounded by difficulties and ethical constraints. The neural tissue is fragile and has scarce regeneration capacity [1], complicating the extraction of primary neural cells without damaging the local tissue and leaving functional sequelae. Therefore, it is essential to the field that cell culture models are developed, and naturally that they are improved to better simulate real neural tissues.

The traditional approach of cell culture has many problems. Firstly, it relies on cell lineages, that have already been passaged for several generations and are (epi)genetically deviated. These cells are mainly transformed, either for being extracted from neoplasms or by artificial mechanisms. The alternative of primary cells exists, but is not always applicable, as in the case of the nervous system. Cells are typically cultured in two-dimensional artificial environments, which cannot reproduce cell-cell and cell-matrix interactions that occur *in vivo* [2]. These interactions are fundamental for tissue characterization. The extracellular matrix (ECM) is a pivotal hub of signaling, essential for several processes, such as cell differentiation, migration, proliferation and morphology [3]. Cell-cell interactions are also essential and ubiquitous, participating in development, cell function, growth and homeostasis, to name just a few [4]. Alterations in both ECM and cell-cell interactions are involved in pathogenic processes [3–5].

In the neural field, the signaling promoted by the microenvironment is also fundamental. It has been described its importance in differentiation [6], neurogenesis [6] and in disease [7]. The vast array of ECM glycoproteins present in mammals influences virtually all aspects of nervous system development and function [8]. Thus, for neuroscience research to remain relevant, we need culture models that simulate the microenvironments and interactions found *in vivo*, a position best filled through three-dimensional models.

Human iPSC-derived neurospheres are three-dimensional aggregates primarily composed of neural stem cells, radial glia-like cells, neural progenitors, and early postmitotic neurons. Under defined differentiation conditions, they can give rise to neurons and astrocytes, recapitulating aspects of early human neurodevelopment in vitro. They are formed from NSCs or progenitor cells cultured in media tipically supplemented with N2 or B27 and growth factors such as bFGF or EGF [9]. The suspended culture generates a cell cluster, which provides a 3D environment for cells to interact with one another and with the ECM they secrete. As previously described [10], the ECM of neurospheres has a neural signature. By combining a 3D organization with cell type heterogeneity and a neural-like ECM, neurospheres can provide the cell interactions and signaling microenvironments that are not well represented in 2D cultures. We chose neurospheres over cerebral organoids because, although organoids offer higher structural and cellular complexity, they also exhibit greater intra- and inter-organoid heterogeneity and high batch-to-batch variability, which can hinder robust comparative proteomic analyses [11,12]. Previous studies have shown that neurospheres provide a standardized and reproducible in vitro model for medium-to high-throughput molecular assays [11,13].

The research applications for neurospheres are diverse, ranging from diseases [14] to development [15], differentiation [16] and cell interactions [17] in the nervous system. Moreover, patient-derived neurospheres are a promising model for personalized medicine. Since they can be made from patients’ cells or iPSCs, individual particularities such as the genetic background can be incorporated into this cell culture model, allowing for the investigation of the impact of personal features on the effectiveness of therapeutic strategies and drugs.

Here, we conducted a proteomics experiment to compare 2D cultured NSCs (our “NSC” samples) with 3D cultured NSCs (forming neurospheres, our “NPH” samples), using hiPSC-derived NSCs. In addition to the proper proteomic characterization of the groups, we implemented a fast experimental methodology, with little material and no offline fractionation. Currently, studies of this kind are relevant to the development of single-cell proteomics, personalized medicine and screening assays, situations that demand speed, scaling or in which fractionation is unsuitable. So, we asked how much depth and relevance could we achieve in a fast analysis, using TMT-based quantification, which allows multiplexing and simultaneous analysis of samples, increasing sensitivity, reproducibility and decreasing time consume [18].

## Materials and Methods

### Samples preparation and culture

Human induced pluripotent stem cells (iPSCs) were sourced from the Coriell Institute for Medical Research repository (GM23279A). The iPSCs were mantained in Stem Flex media (Thermo Fisher Scientific), cultivated on Matrigel-coated (BD Biosciences) dishes at a temperature of 37°C within a humidified atmosphere containing 5% CO_2_. Media was changed regularly, to prevent spontaneous differentiation and colonies were manually split at 70–80% confluence.

For differentiation of iPSCs into NSCs, a PSC neural Induction Medium, consisting of Neurobasal medium and a neural induction supplement (Thermo Fisher Scientific), was used according to the manufacturer’s instructions. Media was renewed every other day until day 7, after which NSCs were separated and expanded in a 1:1 mixture of Advanced DMEM/F12 and Neurobasal medium supplemented with a neural induction supplement (Thermo Fisher Scientific).

For proteomic analysis, neurospheres were formed by suspending NSCs after they reached 90% confluence and were detached using accutase (Merck Millipore), followed by centrifugation at 300 g for 5 minutes. The cells were then resuspended in neural induction medium, consisting of a 1:1 mixture of DMEM/F12 (Life Technologies) and Neurobasal medium (Thermo Fisher Scientific), supplemented with 1× N2 (Invitrogen) and 1× B27 (Thermo Fisher Scientific). Subsequently, 8 × 10^4^ NSCs were placed in each well of a 96-well plate and cultured on an orbital shaker set at 90 rpm for 3h at 37°C in an atmosphere containing 5% CO_2_. In total, 10 samples were made: 5 from 2D cultured neural stem cells (NSCs) and 5 from 3D cultured neural stem cells, forming neurospheres (NPHs).

### Sample processing

The protocol was based on previous studies from our laboratory [19,20]. Samples were washed once with PBS 1×. Then, lysis buffer (8M Urea, 50mM HEPES, pH 8.5) was added. Cells were sonicated at 40 kHz for 10 min. The lysate was centrifuged at 13000 rpm for 10 min, to collect extracted proteins from the supernatant.

Proteins were reduced with dithiothreitol (10mM at 30°C for 1h) and alkylated with iodoacetamide (40mM, room temperature, in the dark for 30 min). The protein concentration was determined by Qubit 4 Fluorometer (Invitrogen), following manufacturer’s protocol. Samples were diluted 10× with 50mM HEPES pH 8.5 buffer, to minimize urea interference with trypsin, and then digested with trypsin (Promega, USA) at a 1:50 enzyme-to-protein ratio overnight at 37°C and 200 rpm. The reaction was quenched with 1% trifluoroacetic acid (TFA).

Samples were dessalted with home-made reversed-phase columns containing C18 matrix and Poros™ 20 R2 resin, using 0.1% TFA as the wash solvent, and 50% or 70% acetonitrile in 0.1% TFA as the elution solvents. After desalting, eluates were vacuum-dried and stored at -80°C.

### Isobaric labeling

Digested peptides were labeled using TMT 10plex™ (Thermo Scientific). Briefly, peptides were resuspended in 50mM HEPES, pH 8.5, and incubated with 13.5mM TMT reagents at room temperature for 1h. Each sample was labeled with a distinct TMT channel (e.g., 126 for NPH_1, 127N for NSC_1, 127C for NPH_2, 128N for NSC_2, etc.). We then pooled 0.8 μg of labeled peptides from each sample (total of 8 μg) in a single tube containing 50 μL of 0.1% TFA. The pooled sample was washed and dried according to the desalting protocol above.

### LC-MS/MS measurements

We performed single-shot LC-MS/MS on an EASY-nLC™ 1000 system coupled to a Q Exactive™ Plus mass spectrometer (Thermo Scientific). After reconstitution in 0.1% TFA, 500 ng of peptides were injected onto a 75-µm × 75-cm EASY-Spray analytical column (Thermo Scientific) packed with C18 resin, at a flow rate of 300 nL/min. The mobile-phase gradient began at 2% solvent B (90% ACN, 0.1% Formic acid) in solvent A (0.1% Formic acid) and proceeded until 95% B in a total of 68 minutes of acquisition. The mass spectrometer was operated in data-dependent acquisition (DDA) mode with positive ionization. MS1 scans were acquired at a resolution of 70k, with an AGC target of 3 × 10^6^ and a maximum injection time (maxIT) of 100 ms. MS2 scans were acquired at a resolution of 35k, with an AGC target of 10^5^ and a maxIT of 100 ms. The cycle time was set to 2 seconds. The pool was analyzed in two technical replicates.

### Data processing and analysis

MS data were searched using Proteome Discoverer software [21] (Thermo Scientific, v3.1.1.93), with Sequest HT search engine queried against Swiss-Prot human revised proteome from UniProt (2023). The enzyme set was trypsin, with maximum of two missed cleavages. We set precursor mass tolerance at 10 ppm and fragment mass tolerance at 0.02 Da. Carbamidomethylation of cysteines was considered a fixed modification, while oxidation of methionine, phosphorylation (S/T/Y) and N-terminal acetylation were set as variable modifications. Cross-referenced TMT was performed following the product data sheet.

The resulting data were imported into Perseus software [22] (v2.0.11) for statistical analysis. Corresponding TMT channels from both technical replicates were averaged. Log_2_ transformations were applied and values were median-subtracted for normalization. We performed Student’s T-test with a p-value threshold of 0.01. We did not use a fold change threshold due to the ratio compression inherent in isobaric labeling [18] (instead, we applied a strict *p*-value cutoff of 0.01). Differentially abundant proteins were explored with the online tools DAVID [23,24] and EnrichR [25–27], for functional enrichment analysis. We used the databases Gene Ontology [28,29], Kegg [30–32], Cell Marker [33] 2024 and PangaoDB [34] Augmented 2021, using our identified proteins set as the background. We used an FDR approach (Benjamini–Hochberg) [35,36] for significance, reported in DAVID’s charts as “Benjamini” and in EnrichR’s ones as “Adjusted P-value”. Genes associated with enrichment annotations can be checked in the tables in Supplementary Material.

Enriched terms with FDR ≤ 0.05 were either fully graphed or manually selected, with priority for neural-related ones. For GO cellular components enrichment, we analyzed the proportion of gene counts of neural-related cellular components out of the total of gene counts from all cellular components enriched (FDR ≤ 0.05), neural or not. We applied a two-proportion Z-test to assess the statistical significance of the change in this proportion between groups, with the R function “prop.test”. The ECM analysis in **Figure 4** was performed with the R package MatrisomeAnalyzeR [37]. Principal component analysis, Pearson correlation heatmap, Volcano plot, Bubble plots and Matrisome pie charts were generated in R scripts. Bar graphs were produced with GraphPad Prism 8. Office package 2016 (Microsoft) was also used for data handling, tables generation and assembly of figures. In the workflow, we used free images from Scidraw and Servier Medical Art. Links to the source of the figures are available in Supplementary material.

The mass spectrometry proteomics data have been deposited to the ProteomeXchange Consortium via the PRIDE [38] partner repository with the dataset identifier PXD063185 and 10.6019/PXD063185.

## Results

After filtering out contaminants and identifications with FDR above 0.01, a total of 1576 proteins were identified. We also had 5827 peptides and 9435 PSMs. We plotted histograms of the median-normalized abundance values for each sample (Supplementary Material) and no normalization issues were found across samples.

The PCA **(Figure 2a)** did cluster the two groups (NSCs and NPHs), meaning they have particular characteristics, reflected at the proteome level, that are typical from each one and that differ one from the other. Sample NSC_3 positioned far from both groups. Pearson correlation heatmap **(Figure 2b)** revealed a slight shift of correlation coefficients between NSC_3 and other NSCs samples. Despite that, all correlations shown are considered high and the heatmap also revealed a clustering of groups (lighter areas).

**Figure 1.**
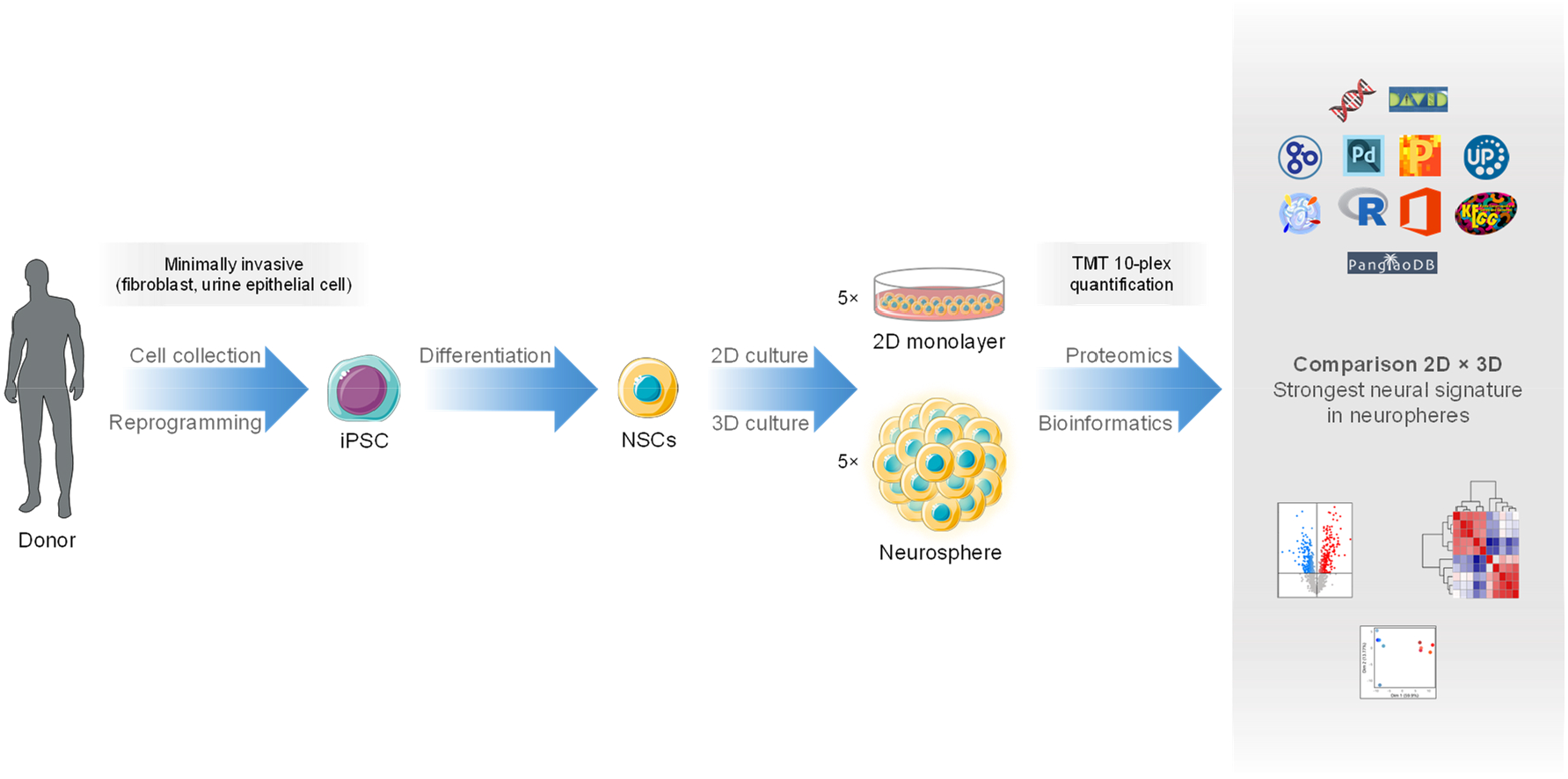
General workflow. iPSCs (GM23279A, Coriell Institute) were differentiated in NSCs, which were cultured in 2D monolayers and 3D neurospheres (five replicates each). Proteins were extracted, digested and the peptides were labeled with the isobaric label TMT 10-plex, enabling multiplexed relative quantification. Pooled peptides were analyzed by LC-MS/MS and several bioinformatics tools were used to analyze the data.

**Figure 2.**
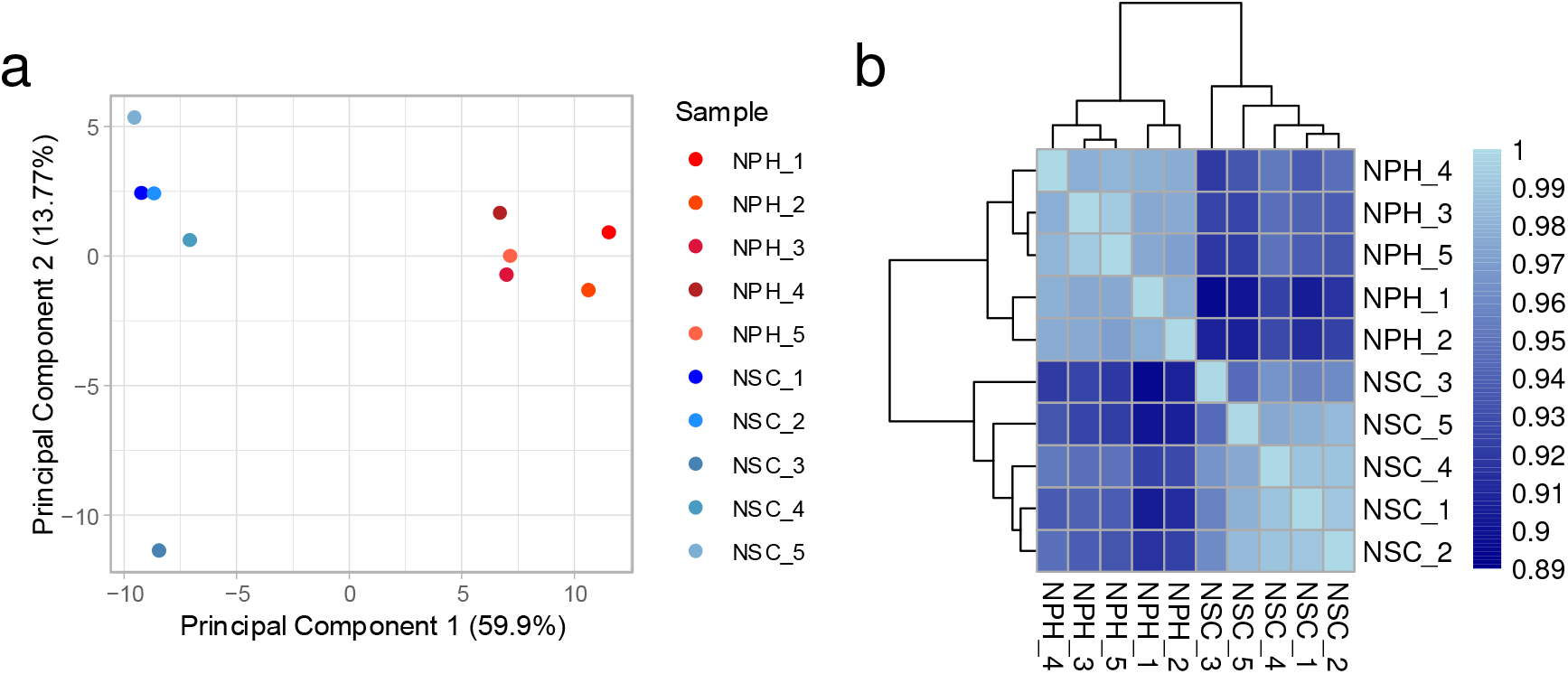
Statistical analysis of group clusterization. **a:** Principal Component Analysis (PCA). 2D NSCs are shown in blue and 3D NSCs (neurospheres) in red. **b:** Heatmap of Pearson correlation coefficients between samples. All samples showed high correlation (r > 0.89). Differences are represented on a dark-to-light blue gradient scale.

We performed Student’s T-test **(Figure 3a)** to assess proteins fold changes between groups and their statistical significance. Using *p*-value cutoff of 0.01 and no fold change cutoff, we obtained 271 proteins more abundant in NSCs (Down NPH/NSC) and 271 proteins more abundant in neurospheres (Up NPH/NSC). We also performed the same analysis but excluding sample NSC_3, which returned 267 proteins Down NPH/NSC and 288 proteins Up NPH/NSC, suggesting that no major difference occurred.

**Figure 3.**
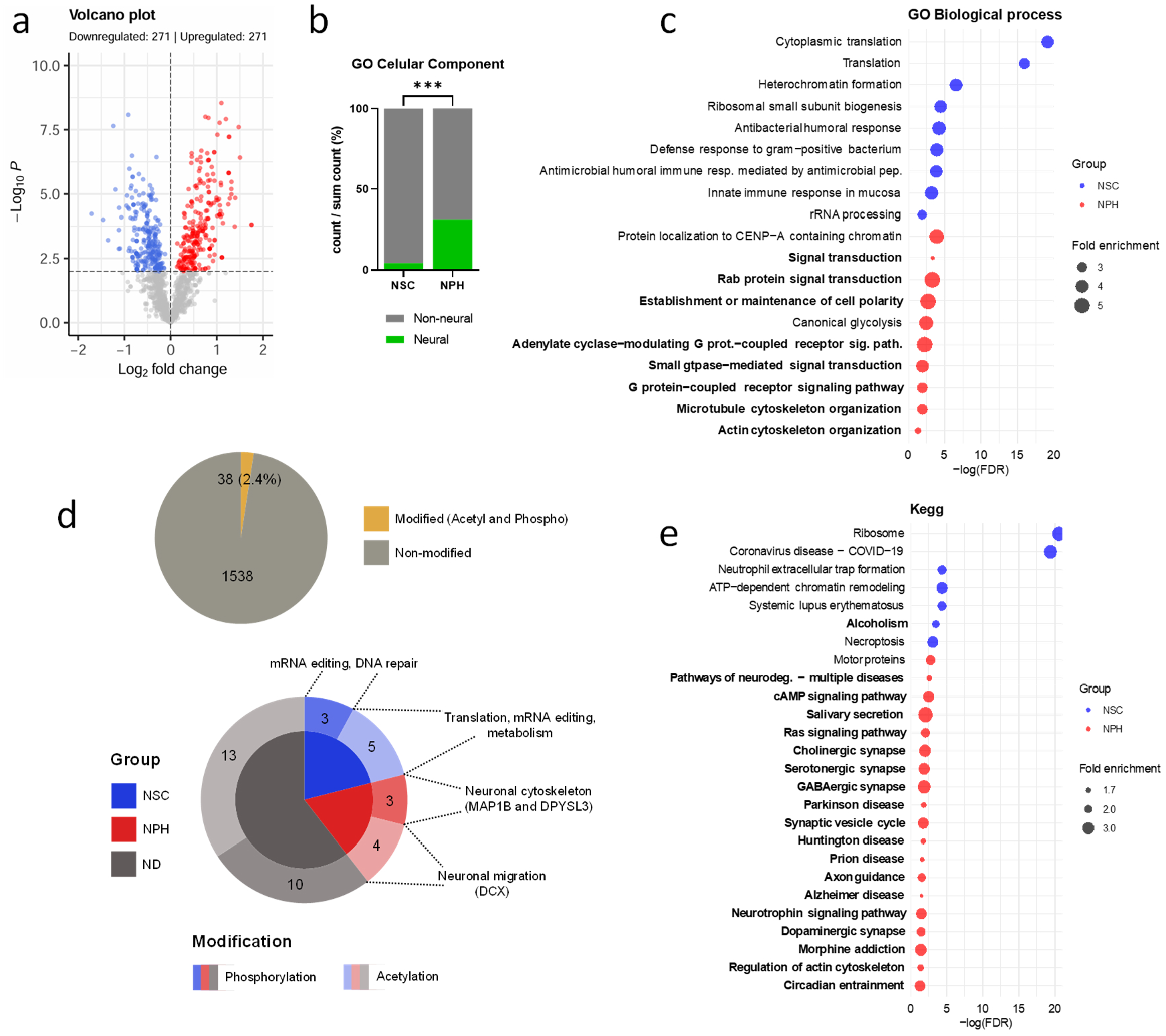
Analysis of differentially abundant proteins by functional enrichment. **a:** Volcano plot showing the results of Student’s T-test. In blue, proteins more abundant in 2D NSCs and in red, proteins more abundant in neurospheres. We used *p*-value cutoff of 0.01 and no fold change cutoff. **b:** Proportion of gene counts of neural-related Gene Ontology enrichment cellular components out of the total gene counts matched to all enriched cellular components (neural and non-neural). ^***^: *p*-value < 0.001. **c:** Bubble plot of Gene Ontology enrichment biological processes. The position of the circles related to the x axis reflects their value of -log(FDR), so higher values indicate more statistical significance. Their size reflects the fold enrichment and the colors, the experimental group. All enriched NSCs terms are shown. Terms of NPHs were selected, with priority to neural-related ones (in bold). Only terms with FDR ≤ 0.05 were considered. **d:** Analysis of post-translational modifications (PTMs). The upper pie chart shows the slice of our dataset which was phosphorylated or acetylated. The lower one shows the distribution of each modification and whether it was differentially abundant. The dotted lines show the physiologic involvement of the proteins from each classification and the numbers represent the number of proteins in each. ND: non-differential. **e:** Bubble plot of Kegg enrichment. All enriched terms of NSCs and all neural-related ones of NPHs are shown. Other parameters are the same as in **b**.

**Figure 4.**
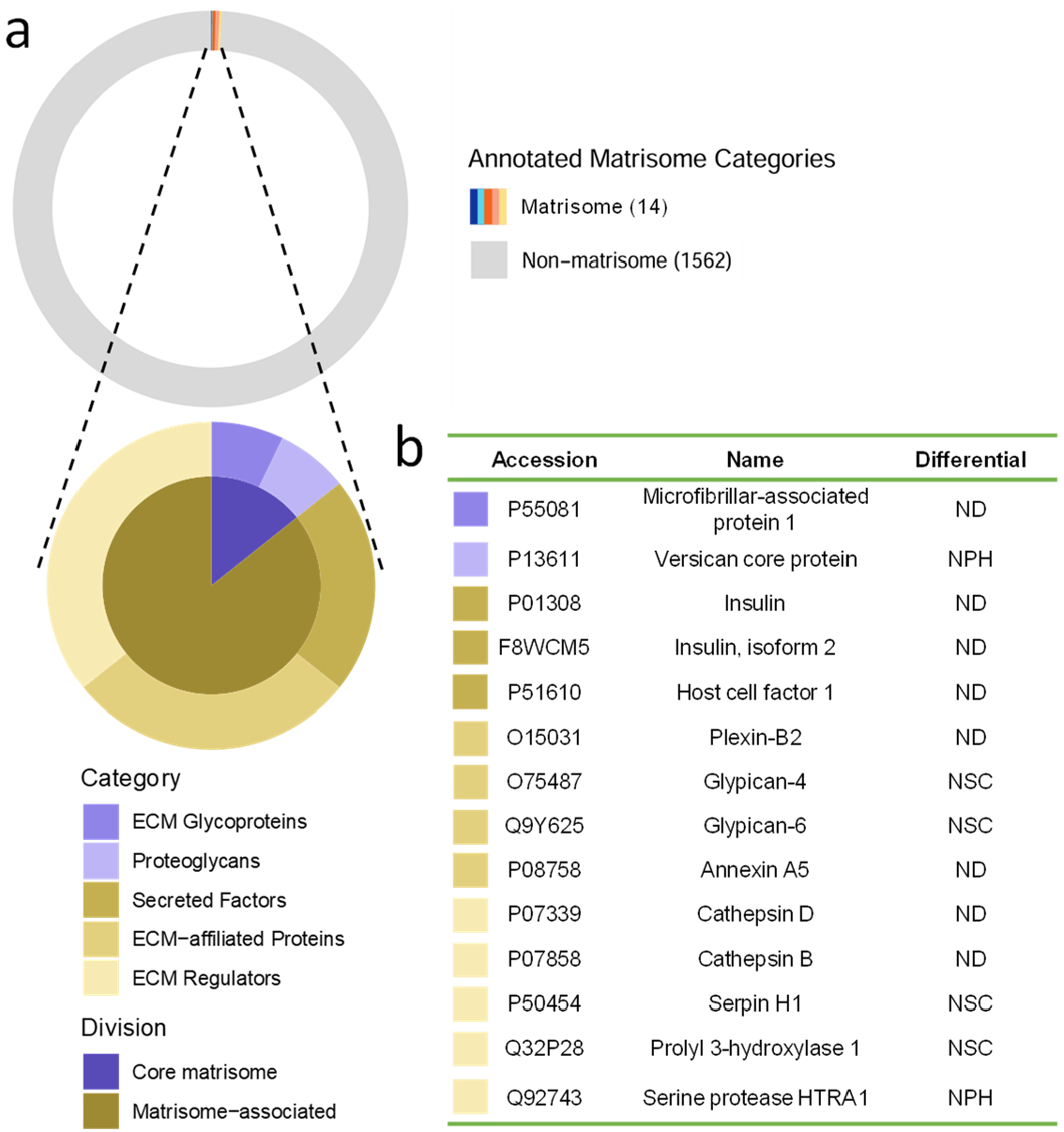
Analysis of Extracellular matrix-related proteins. Proteins were annotaded with MatrisomeAnalyzeR package, which uses the database of the Matrisome Project. **a:** Pie charts showing the slice of our dataset annotaded as ECM-related (“Matrisome”) and the proportion of each ECM category inside this slice. Core matrisome proteins are displayed in purple and matrisome-associated ones, in brown. **b:** Table presenting the 14 proteins associated with the ECM annotaded in **a**. The colors before accessions designate the ECM category to which the protein belongs and follows the same code from the previous legend. The “Differential” column shows if the protein is differentially abundant between groups. ND: non-differential.

Differentially abundant proteins were analyzed with functional enrichment using DAVID web tool. Gene Ontology (GO) analysis for biological processes **(Figure 3c)** revealed no neural-related process enriched in NSCs with FDR ≤ 0.05. The enriched processes were mainly involved with protein translation and immune response. In neurospheres, by contrast, we observed 8 neural-related enriched processes, related to signal transduction/signaling pathways, cell polarity, cytoskeleton organization and Rab signal transduction. Just for scale purposes, “Signal transduction” fold enrichment was of approximately 2.2. Kegg enrichment **(Figure 3e)** annotated just one neural-related pathway enriched in NSCs, which was alcoholism. Enriched processes were diverse, ranging from protein translation to diseases, chromatin remodeling and others. In neurospheres, there were 19 neural-related pathways enriched. They were mainly involved with disease and synapse, but also included axon guidance, neurotrophin signaling pathway and circadian entrainment, for instance.

GO enrichment for cellular components **(Figure 3b)** revealed an increase of approximately 7.3-fold from NSCs to neurospheres in the proportion of gene counts of neural-related cellular components out of the total gene counts of all cellular components enriched, neural and non-neural. We performed a two-proportion Z-test in R to verify the statistical significance about the difference between the proportions. The *p*-value associated with the difference was below 2.2 × 10^−16^, indicating an extremely low chance of this result being random and not due to a real difference between groups.

PTM analysis **(Figure 3d)** showed that just 38 of the identified proteins were phosphorylated or acetylated. Most of the PTMs were not differentially abundant and the quantity of modifications differential in each group differed just by one (NSCs had more). The proteins modified in NSCs were related to mRNA editing, metabolism and DNA repair. The ones of NPHs were, among others, related to neuronal cytoskeleton, involved in processes such as neuron migration, neurite extension and axon guidance (those are indicated by gene name in parentheses).

We analyzed differentially abundant proteins without sample NSC_3 and achieved almost the same results. GO enrichment for biological processes revealed a slight decrease in enrichment significance overall, Kegg enrichment was almost identical and GO enrichment for cellular components was identical. We concluded that these minor differences did not justify the exclusion of sample NSC_3 from the analysis. Biological variability between biological replicates are expected [19] and likely account for the slight differences seen with NSC_3.

We performed a search to annotate the proteins in our dataset which are related to ECM using the R package MatrisomeAnalyzeR, which searches against the dataset of the Matrisome Project **(Figure 4)**. Only 14 proteins were annotated as ECM-related, most of them matrisome-associated. The division “Core matrisome” refers to structural ECM proteins and “Matrisome-associated” refers to non-structural ones, including proteins that structurally resemble core matrisome proteins (“ECM-affiliated”), regulate and modify the ECM (“ECM Regulators”) and are signaling molecules (“Secreted Factors”). Four of the annotated proteins had increased abundance in NSCs and two in NPHs (including one core matrisome protein).

We then searched for cell type markers and enrichments to understand the composition of each model. First, we searched manually for cell type markers of undifferentiated and differentiated neural cells **(Table 1)**. The markers SOX2 and NES, which identify undifferentiated neural cells, had increased abundance in NSCs. The markers MAP2, TUBB3 and neurofilament proteins Light, Medium and Heavy characterize differentiated neurons and had increased abundance in NPHs. The other identified markers were not differentially abundant. The only searched marker of astrocytes identified was GFAP. None of the searched oligodendrocytes markers were identified. Cell type enrichments were performed through EnrichR web tool. Cell Marker 2024 database enrichment **(Figure 5a)** returned one neural-related term for NSCs, “Microglial Cell Brain Human”, and four for NPHs (neural-related terms are in bold). NSCs enriched mainly for non-neural cell types. In PanglaoDB Augmented 2021 database enrichment **(Figure 5b)**, NSCs enriched just for “Pluripotent Stem Cells” and NPHs enriched for several neural cell types. To see the genes matched to each annotation please check Supplementary Material.

**Table 1.**
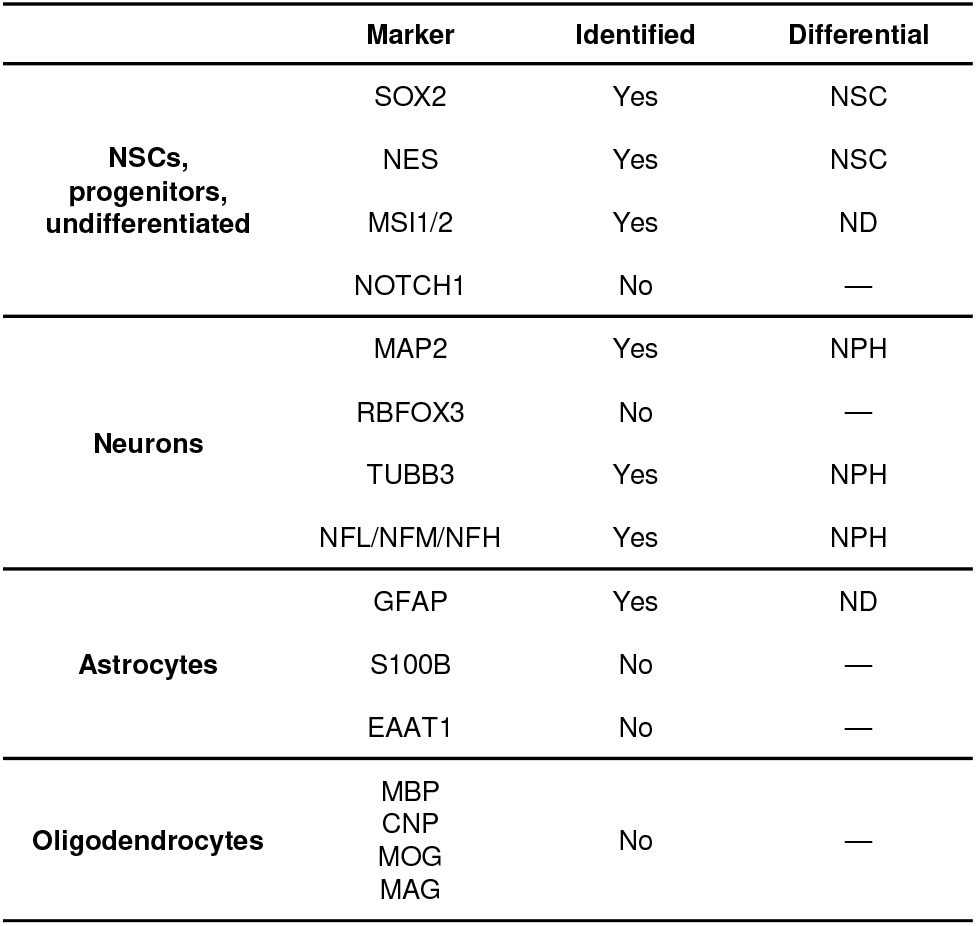
Search of cell type markers [39–42]. Markers are displayed by gene names. The “Differential” column shows if the protein is differentially abundant between groups. ND: non-differential. —: not applicable.

**Figure 5.**
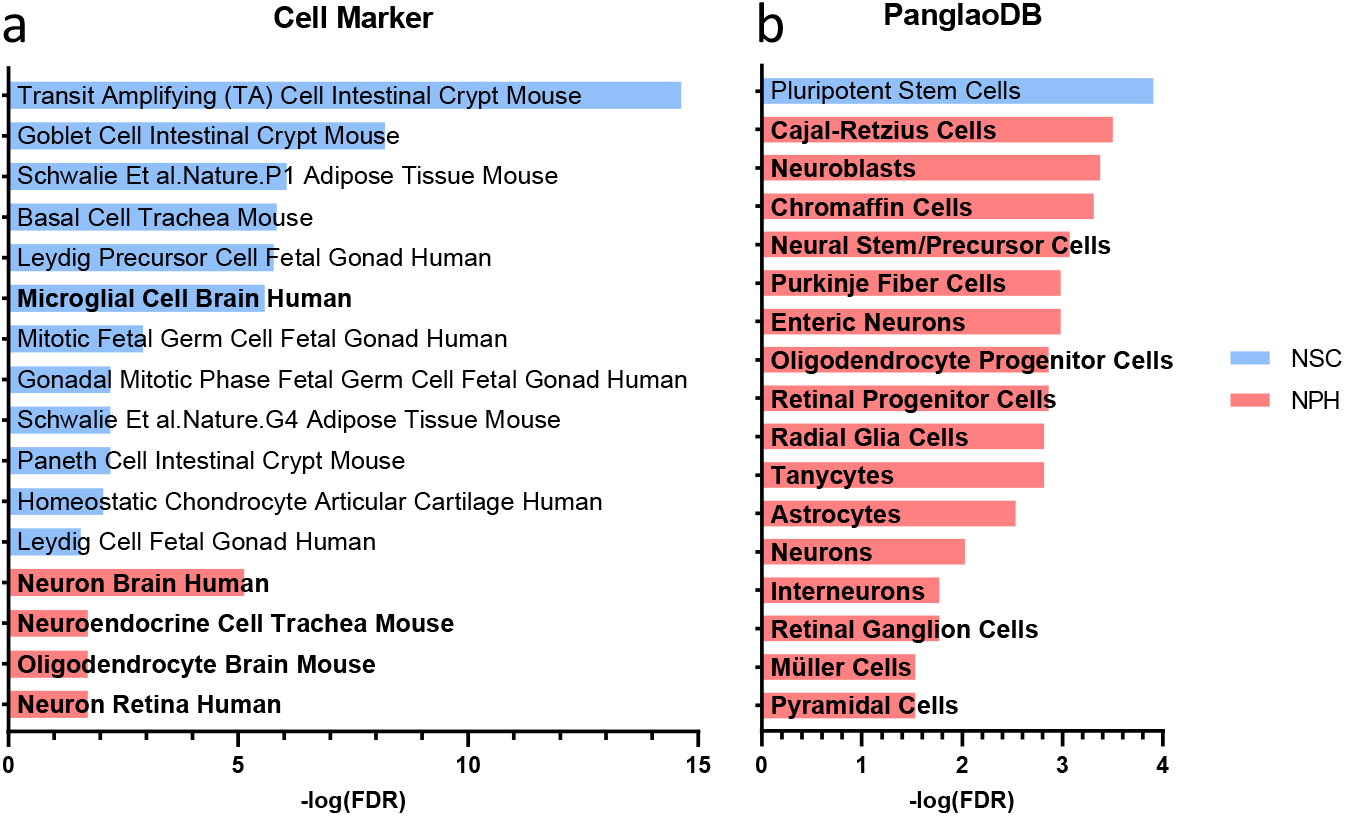
Analysis of cell type enrichment. **a:** Enrichment of cell type using Cell Marker 2024 database from EnrichR web tool. Annotations are graphed by their values of - log(FDR), which increases as the statistical significance increases. All annotations with FDR ≤ 0.05 are displayed. Neural-related terms are in bold. **b:** Enrichment of cell type using PanglaoDB Augmented 2021 database from EnrichR web tool. NSCs had only one annotation and annotations from NPHs were manually selected (all with FDR ≤ 0.05). Other parameters are the same as in **a**.

We asked how much of a relevant pathway we could cover. So we mapped the identified proteins from the Kegg pathway “Synaptic Vesicle Cycle” (Supplementary Material). Result showed that more than a half of the pathway could not be identified. Of the identified proteins, almost all had increased abundance in NPHs and one was not differential.

## Discussion

Our results show a great proteomic difference between 2D- and 3D-cultured NSCs. Out of the 1576 proteins identified 542 were differentially abundant between the groups. Underscoring this, with a *p*-value cutoff of 0.01 and fold change cutoff of 2^0.5^, we still have 287 differentially abundant proteins (not shown). This is especially impressive given the inherent ratio-compression effect of isobaric label-based quantification (such as TMT), which typically attenuates measured fold changes [18]. PCA and Pearson correlation **(Figure 2)** also separated the groups.

Many of the enriched processes in neurospheres are synapse-related, namely G protein and cAMP signaling, the more general “signal transduction” processes and the many specific synapse processes **(Figure 3)**. The enriched process “Rab protein signal transduction” **(Figure 3c)** is of extreme importance for neural function. Rab proteins are small GTPases that participate in all steps of intracellular vesicle trafficking, from vesicle formation to movement, docking and fusion with the plasma membrane [43,44]. Being the major regulators of vesicle trafficking, they are essential for chemical synapses, in which synaptic vesicles containing neurotransmitters are exocytosed. Specific Rab subtypes, specially Rab3, have long been linked to neurotransmission [43,45]. Neurodegenerative processes are correlated with altered Rab-mediated vesicle trafficking and Rab mutations have been identified in neurodegeneration[46]. We identified Rab proteins related to neural function, such as Rab-3A, Rab-4A, Rab-15 and Rab-39B. We also identified Rab GDP dissociation inhibitor alpha (GDI1), which promotes the deactivation of Rab proteins.

Enrichment in neurospheres of processes related to cytoskeleton and cell polarity **(Figure 3c)** suggests that the cells in this model undergo morphological organization. In neural cells, the cytoskeleton is fundamental for maintaining the anisotropic cell shape and the overall cell function [47]. By anisotropic we mean that the morphology of a neuron is characterized by three distinct regions – dendrites, soma and axon. These different regions vary drastically in shape, and those shapes are physically maintained by the cytoskeleton scaffold. Moreover, the cytoskeleton is responsible for synaptic vesicles transport along the extensive axons, with the help of motor proteins, to the axon terminal, where they fuse with the membrane and release the neurotransmitter content in the synaptic cleft [47]. Interestingly, the process “Motor proteins” was also enriched **(Figure 3e)**.

The neural-related processes enriched in neurospheres are evidence for an enhanced neural differentiation. “Neurotrophin signaling pathway” **(Figure 3e)**, underlies several processes, including differentiation of NSCs into neurons, survival of neural cells (some cases), synaptogenesis and even axon initiation and guidance [48,49]. Axon guidance, the process which directs axonal outgrowth by chemotaxis, was also enriched in our data **(Figure 3e)**. The ability of 2D and 3D neural culture models to generate synaptic connections (or an indication of this) has already been described [50,11,51]. In cell type analysis **(Table 1 and Figure 5)** 2D NSCs had increased abundance of multipotency markers and enriched mainly for non-neural cell types. Meanwhile, NPHs had increased abundance of neuronal markers and enriched for several neural cell types. While 2D NSCs express more general proteins, 3D NSCs express more proteins related to the nervous system, a fact that can be observed in our enrichment analyses.

Many neurological disorders were enriched in neurospheres, such as Parkinson disease and morphine addiction **(Figure 3e)**. This suggests the potential to simulate neurological disorders *in vitro* using 3D models. In fact, this possibility is already being explored, as we now have studies describing the ability of 3D models to recapitulate key pathophysiological characteristics of neurological diseases [14,52–54]. Besides research applications, the establishment of disease-relevant 3D cell culture models will contribute a lot to personalized medicine, enabling *in vitro* assays to explore patients’ genotype and predict therapeutic outcomes. Relevant culture models also reduce the necessity for animal models for research and allow easier, cheaper and less time-consuming scaling.

Our analysis of ECM-related proteins **(Figure 4)** retrieved little results, only 14 proteins. Of those, 4 had increased abundance in NSCs and 2 in NPHs, the opposite that one would think, since the secretion of an ECM is a hallmark of three-dimensional cell culture models. However, the analysis of the ECM with proteomics is surrounded by intrinsic difficulties of these structures [55]. A specialized pipeline to enrich ECM proteins is important and would increase the depth of our identifications and allow a more reliable comparison between groups. We also had small results in PTM analysis **(Figure 3d)**, only 38 proteins were phosphorylated or acetylated. Groups differed just by one in the number of differential modifications and some proteins modified in NPHs were neural-related. Again, we could not achieve sufficient depth without a PTM-focused enrichment.

Our analysis of the coverage of the Kegg pathway “Synaptic Vesicle Cycle” (Supplementary Material) revealed that we could not cover it with much depth. We can clearly see that the differential proteins from NPHs were prevalent, but a real experimental exploration of this pathway may be compromised by the missing identifications. With newer mass spectrometers we would probably have a greater coverage and maybe a real exploration of this pathway would be possible together with a fast workflow.

When non-transformed cells are highly confluent, they undergo a cell cycle arrest, that is called “contact inhibition of proliferation”. This is a consequence of the activation of the Hippo signaling pathway by mechanical signals, which causes phosphorylation of YAP/TAZ and their localization in the cytosol (inactive state) [56]. In consequence, autophagy and proliferation are inhibited. We hypothesized that this would occur in neurospheres, since cells in this model have more physical contact. We identified only the YAP1 protein in our data, which had increased abundance in NSCs (result not shown). However, we cannot determine subcellular localization (active in the nucleus or inactive in the cytosol), since in our protocol we sonicated the samples, leading to rupture of all cell membranes and mixture of their contents. It was not identified phosphorylation of YAP, but we did not perform an enrichment for phosphopeptides, which limits our insights about this pathway.

Our results clearly show an increase in the neural signature of neural stem cells proteome when transitioning from 2D to 3D culture. To explain such a great difference, 3D culture of human neural stem cells likely promoted epigenetic changes that drove the cells to express a proteomic profile more representative of mature neural tissue. Previous studies have reported that 3D *in vitro* neural cell culture models exhibit stronger neural signature than 2D ones, corroborating our findings [10–12,57]. However, a fast method like ours may not be suitable for occasions in which a special enrichment in necessary, such as in ECM and PTM investigation. In single-cell analysis, a more specialized workflow is preferred [58]. Low-abundance protein analysis, which is enhanced by offline fractionation, longer runs and other strategies, such as high-abundance protein depletion, can also be considered a weak point of our approach. However, modern mass spectrometers would certainly increase the depth of our analysis and could possibly overcome these limitations.

In all analyses presented here, neurospheres of iPSC-derived NSCs demonstrated to be closer to a real neural tissue than monolayers of the same cells in the proteome level. With them, we have a more reliable and predictive model for research and personalized medicine, expanding the possibilities of neural modeling while reducing dependency on animal models and primary cultures.

## Supporting information

Table of cell type enrichments

Table of GO biological process enrichment

Table of GO celular component enrichment

Table of Kegg enrichment

Table with identified proteins

## Author contributions

**Pedro de Lima Muniz:** Writing - Original Draft, Project administration, Methodology, Investigation, Formal analysis. **Leticia Rocha Quintino Souza:** Methodology (Samples preparation and culture). **Michele Rodrigues Martins:** Writing - Review & Editing, Validation. **Wassali Valadares:** Methodology (LC-MS/MS measurements). **Fábio César Sousa Nogueira:** Writing - Review & Editing, Resources. **Stevens Rehen:** Writing - Review & Editing, Resources. **Guillaume Nugue:** Writing – Original Draft, Supervision, Conceptualization, Methodology. **Magno Junqueira:** Writing - Original Draft, Supervision, Conceptualization, Funding acquisition.

All authors have read, reviewed and approved this manuscript.

## Funding

This work was funded by Fundação Carlos Chagas Filho de Amparo à Pesquisa do Estado do Rio de Janeiro (FAPERJ, project APQ1-FAPERJ 313417/2021-0) and Conselho Nacional de Desenvolvimento Científico e Tecnológico (CNPq). The funding agencies had no involvement in any step of the research. Pedro de Lima Muniz received a PIBIC-UFRJ scholarship for this project.

## Declaration of competing interest

The authors declare that they have no known competing financial interests or personal relationships that could have appeared to influence the work reported in this paper.

## Declaration of generative AI and AI-assisted technologies in the writing process

During the preparation of this work the author(s) used ChatGPT (OpenAI) in order to improve sentences. After using this tool/service, the author(s) reviewed and edited the content as needed and take(s) full responsibility for the content of the publication.

## Acknowledgements

Our sincere thanks to the groups of the professors Stevens Rehen, Fábio César Sousa Nogueira and Magno Junqueira, for the fruitful discussions. We also sincerely thank the Federal University of Rio de Janeiro, the public research funding agencies FAPERJ (project APQ1-FAPERJ 313417/2021-0) and CNPq and the teams of all the tools we used.

## Notes

### Competing Interest Statement

The authors have declared no competing interest.

## References

[1] A. Carnicer-Lombarte, D.G. Barone, F. Wronowski, G.G. Malliaras, K. Franze, Regenerative capacity of neural tissue scales with changes in tissue mechanics post injury, Biomaterials 303 (2023) 122393. 10.1016/j.biomaterials.2023.122393.

[2] P. Horvath, N. Aulner, M. Bickle, A.M. Davies, E.D. Nery, D. Ebner, M.C. Montoya, P. Östling, V. Pietiäinen, L.S. Price, S.L. Shorte, G. Turcatti, C. von Schantz, N.O. Carragher, Screening out irrelevant cell-based models of disease, Nat. Rev. Drug Discov. 15 (2016) 751–769. 10.1038/nrd.2016.175.

[3] N.K. Karamanos, A.D. Theocharis, Z. Piperigkou, D. Manou, A. Passi, S.S. Skandalis, D.H. Vynios, V. Orian-Rousseau, S. Ricard-Blum, C.E.H. Schmelzer, L. Duca, M. Durbeej, N.A. Afratis, L. Troeberg, M. Franchi, V. Masola, M. Onisto, A guide to the composition and functions of the extracellular matrix, FEBS J. 288 (2021) 6850–6912. 10.1111/febs.15776.

[4] J. Su, Y. Song, Z. Zhu, X. Huang, J. Fan, J. Qiao, F. Mao, Cell–cell communication: new insights and clinical implications, Signal Transduct. Target. Ther. 9 (2024) 1– 52. 10.1038/s41392-024-01888-z.

[5] L. Peng, F. Wang, Z. Wang, J. Tan, L. Huang, X. Tian, G. Liu, L. Zhou, Cell–cell communication inference and analysis in the tumour microenvironments from single-cell transcriptomics: data resources and computational strategies, Brief. Bioinform. 23 (2022) bbac234. 10.1093/bib/bbac234.

[6] J.-C. Xu, M.-F. Xiao, I. Jakovcevski, E. Sivukhina, G. Hargus, Y.-F. Cui, A. Irintchev, M. Schachner, C. Bernreuther, The extracellular matrix glycoprotein tenascin-R regulates neurogenesis during development and in the adult dentate gyrus of mice, J. Cell Sci. 127 (2014) 641–652. 10.1242/jcs.137612.

[7] I. Gregorio, P. Braghetta, P. Bonaldo, M. Cescon, Collagen VI in healthy and diseased nervous system, Dis. Model. Mech. 11 (2018) dmm032946. 10.1242/dmm.032946.

[8] C.S. Barros, S.J. Franco, U. Müller, Extracellular Matrix: Functions in the Nervous System, Cold Spring Harb. Perspect. Biol. 3 (2011) a005108. 10.1101/cshperspect.a005108.

[9] J.B. Jensen, M. Parmar, Strengths and limitations of the neurosphere culture system, Mol. Neurobiol. 34 (2006) 153–161. 10.1385/MN:34:3:153.

[10] D. Simão, M.M. Silva, A.P. Terrasso, F. Arez, M.F.Q. Sousa, N.Z. Mehrjardi, T. Šarić, P. Gomes-Alves, N. Raimundo, P.M. Alves, C. Brito, Recapitulation of Human Neural Microenvironment Signatures in iPSC-Derived NPC 3D Differentiation, Stem Cell Rep. 11 (2018) 552–564. 10.1016/j.stemcr.2018.06.020.

[11] P. Zhuang, A.X. Sun, J. An, C.K. Chua, S.Y. Chew, 3D neural tissue models: From spheroids to bioprinting, Biomaterials 154 (2018) 113–133. 10.1016/j.biomaterials.2017.10.002.

[12] S. Logan, T. Arzua, S.G. Canfield, E.R. Seminary, S.L. Sison, A.D. Ebert, X. Bai, Studying Human Neurological Disorders Using Induced Pluripotent Stem Cells: from 2D Monolayer to 3D Organoid and Blood Brain Barrier Models, Compr. Physiol. 9 (2019) 565. 10.1002/cphy.c180025.

[13] A.P. Terrasso, C. Pinto, M. Serra, A. Filipe, S. Almeida, A.L. Ferreira, P. Pedroso, C. Brito, P.M. Alves, Novel scalable 3D cell based model for in vitro neurotoxicity testing: Combining human differentiated neurospheres with gene expression and functional endpoints, J. Biotechnol. 205 (2015) 82–92. 10.1016/j.jbiotec.2014.12.011.

[14] W.K. Raja, E. Neves, C. Burke, X. Jiang, P. Xu, K.J. Rhodes, V. Khurana, R.H. Scannevin, C.Y. Chung, Patient-derived three-dimensional cortical neurospheres to model Parkinson’s disease, PLOS ONE 17 (2022) e0277532. 10.1371/journal.pone.0277532.

[15] P.P. Garcez, E.C. Loiola, R. Madeiro da Costa, L.M. Higa, P. Trindade, R. Delvecchio, J.M. Nascimento, R. Brindeiro, A. Tanuri, S.K. Rehen, Zika virus impairs growth in human neurospheres and brain organoids, Science 352 (2016) 816–818. 10.1126/science.aaf6116.

[16] T. Gonmanee, T. Arayapisit, K. Vongsavan, C. Phruksaniyom, H. Sritanaudomchai, Optimal culture conditions for neurosphere formation and neuronal differentiation from human dental pulp stem cells, J. Appl. Oral Sci. 29 (2021) e20210296. 10.1590/1678-7757-2021-0296.

[17] F. Pampaloni, E.G. Reynaud, E.H.K. Stelzer, The third dimension bridges the gap between cell culture and live tissue, Nat. Rev. Mol. Cell Biol. 8 (2007) 839–845. 10.1038/nrm2236.

[18] N. Rauniyar, J.R.I. Yates, Isobaric Labeling-Based Relative Quantification in Shotgun Proteomics, J. Proteome Res. 13 (2014) 5293–5309. 10.1021/pr500880b.

[19] J.R. Murillo, I. Pla, L. Goto-Silva, F.C.S. Nogueira, G.B. Domont, Y. Perez-Riverol, A. Sánchez, M. Junqueira, Mass spectrometry evaluation of a neuroblastoma SH-SY5Y cell culture protocol, Anal. Biochem. 559 (2018) 51–54. 10.1016/j.ab.2018.08.013.

[20] G. Nugue, M. Martins, G. Vitória, B.L.D.M.L. Guimaraes, M. Quiñones-Vega, S. Rehen, M.Z. Guimarães, M. Junqueira, Optimized pipeline for personalized neurobiological insights from single patient-derived Neurospheres, J. Proteomics 313 (2025) 105368. 10.1016/j.jprot.2024.105368.

[21] B.C. Orsburn, Proteome Discoverer—A Community Enhanced Data Processing Suite for Protein Informatics, Proteomes 9 (2021) 15. 10.3390/proteomes9010015.

[22] S. Tyanova, T. Temu, P. Sinitcyn, A. Carlson, M.Y. Hein, T. Geiger, M. Mann, J. Cox, The Perseus computational platform for comprehensive analysis of (prote)omics data, Nat. Methods 13 (2016) 731–740. 10.1038/nmeth.3901.

[23] D.W. Huang, B.T. Sherman, R.A. Lempicki, Systematic and integrative analysis of large gene lists using DAVID bioinformatics resources, Nat. Protoc. 4 (2009) 44– 57. 10.1038/nprot.2008.211.

[24] B.T. Sherman, M. Hao, J. Qiu, X. Jiao, M.W. Baseler, H.C. Lane, T. Imamichi, W. Chang, DAVID: a web server for functional enrichment analysis and functional annotation of gene lists (2021 update), Nucleic Acids Res. 50 (2022) W216–W221. 10.1093/nar/gkac194.

[25] E.Y. Chen, C.M. Tan, Y. Kou, Q. Duan, Z. Wang, G.V. Meirelles, N.R. Clark, A. Ma’ayan, Enrichr: interactive and collaborative HTML5 gene list enrichment analysis tool, BMC Bioinformatics 14 (2013) 128. 10.1186/1471-2105-14-128.

[26] M.V. Kuleshov, M.R. Jones, A.D. Rouillard, N.F. Fernandez, Q. Duan, Z. Wang, S. Koplev, S.L. Jenkins, K.M. Jagodnik, A. Lachmann, M.G. McDermott, C.D. Monteiro, G.W. Gundersen, A. Ma’ayan, Enrichr: a comprehensive gene set enrichment analysis web server 2016 update, Nucleic Acids Res. 44 (2016) W90–97. 10.1093/nar/gkw377.

[27] Z. Xie, A. Bailey, M.V. Kuleshov, D.J.B. Clarke, J.E. Evangelista, S.L. Jenkins, A. Lachmann, M.L. Wojciechowicz, E. Kropiwnicki, K.M. Jagodnik, M. Jeon, A. Ma’ayan, Gene Set Knowledge Discovery with Enrichr, Curr. Protoc. 1 (2021) e90. 10.1002/cpz1.90.

[28] M. Ashburner, C.A. Ball, J.A. Blake, D. Botstein, H. Butler, J.M. Cherry, A.P. Davis, K. Dolinski, S.S. Dwight, J.T. Eppig, M.A. Harris, D.P. Hill, L. Issel-Tarver, A. Kasarskis, S. Lewis, J.C. Matese, J.E. Richardson, M. Ringwald, G.M. Rubin, G. Sherlock, Gene Ontology: tool for the unification of biology, Nat. Genet. 25 (2000) 25–29. 10.1038/75556.

[29] The Gene Ontology Consortium, S.A. Aleksander, J. Balhoff, S. Carbon, J.M. Cherry, H.J. Drabkin, D. Ebert, M. Feuermann, P. Gaudet, N.L. Harris, D.P. Hill, R. Lee, H. Mi, S. Moxon, C.J. Mungall, A. Muruganugan, T. Mushayahama, P.W. Sternberg, P.D. Thomas, K. Van Auken, J. Ramsey, D.A. Siegele, R.L. Chisholm, P. Fey, M.C. Aspromonte, M.V. Nugnes, F. Quaglia, S. Tosatto, M. Giglio, S. Nadendla, G. Antonazzo, H. Attrill, G. dos Santos, S. Marygold, V. Strelets, C.J. Tabone, J. Thurmond, P. Zhou, S.H. Ahmed, P. Asanitthong, D. Luna Buitrago, M.N. Erdol, M.C. Gage, M. Ali Kadhum, K.Y.C. Li, M. Long, A. Michalak, A. Pesala, A. Pritazahra, S.C.C. Saverimuttu, R. Su, K.E. Thurlow, R.C. Lovering, C. Logie, S. Oliferenko, J. Blake, K. Christie, L. Corbani, M.E. Dolan, H.J. Drabkin, D.P. Hill, L. Ni, D. Sitnikov, C. Smith, A. Cuzick, J. Seager, L. Cooper, J. Elser, P. Jaiswal, P. Gupta, P. Jaiswal, S. Naithani, M. Lera-Ramirez, K. Rutherford, V. Wood, J.L. De Pons, M.R. Dwinell, G.T. Hayman, M.L. Kaldunski, A.E. Kwitek, S.J.F. Laulederkind, M.A. Tutaj, M. Vedi, S.-J. Wang, P. D’Eustachio, L. Aimo, K. Axelsen, A. Bridge, N. Hyka-Nouspikel, A. Morgat, S.A. Aleksander, J.M. Cherry, S.R. Engel, K. Karra, S.R. Miyasato, R.S. Nash, M.S. Skrzypek, S. Weng, E.D. Wong, E. Bakker, T.Z. Berardini, L. Reiser, A. Auchincloss, K. Axelsen, G. Argoud-Puy, M.-C. Blatter, E. Boutet, L. Breuza, A. Bridge, C. Casals-Casas, E. Coudert, A. Estreicher, M. Livia Famiglietti, M. Feuermann, A. Gos, N. Gruaz-Gumowski, C. Hulo, N. Hyka-Nouspikel, F. Jungo, P. Le Mercier, D. Lieberherr, P. Masson, A. Morgat, I. Pedruzzi, L. Pourcel, S. Poux, C. Rivoire, S. Sundaram, A. Bateman, E. Bowler-Barnett, H. Bye-A-Jee, P. Denny, A. Ignatchenko, R. Ishtiaq, A. Lock, Y. Lussi, M. Magrane, M.J. Martin, S. Orchard, P. Raposo, E. Speretta, N. Tyagi, K. Warner, R. Zaru, A.D. Diehl, R. Lee, J. Chan, S. Diamantakis, D. Raciti, M. Zarowiecki, M. Fisher, C. James-Zorn, V. Ponferrada, A. Zorn, S. Ramachandran, L. Ruzicka, M. Westerfield, The Gene Ontology knowledgebase in 2023, Genetics 224 (2023) iyad031. 10.1093/genetics/iyad031.

[30] M. Kanehisa, S. Goto, KEGG: Kyoto Encyclopedia of Genes and Genomes, Nucleic Acids Res. 28 (2000) 27–30. 10.1093/nar/28.1.27.

[31] M. Kanehisa, Toward understanding the origin and evolution of cellular organisms, Protein Sci. 28 (2019) 1947–1951. 10.1002/pro.3715.

[32] M. Kanehisa, M. Furumichi, Y. Sato, Y. Matsuura, M. Ishiguro-Watanabe, KEGG: biological systems database as a model of the real world, Nucleic Acids Res. 53 (2025) D672–D677. 10.1093/nar/gkae909.

[33] C. Hu, T. Li, Y. Xu, X. Zhang, F. Li, J. Bai, J. Chen, W. Jiang, K. Yang, Q. Ou, X. Li, P. Wang, Y. Zhang, CellMarker 2.0: an updated database of manually curated cell markers in human/mouse and web tools based on scRNA-seq data, Nucleic Acids Res. 51 (2023) D870–D876. 10.1093/nar/gkac947.

[34] O. Franzén, L.-M. Gan, J.L.M. Björkegren, PanglaoDB: a web server for exploration of mouse and human single-cell RNA sequencing data, Database 2019 (2019) baz046. 10.1093/database/baz046.

[35] Y. Benjamini, Y. Hochberg, Controlling the False Discovery Rate: A Practical and Powerful Approach to Multiple Testing, J. R. Stat. Soc. Ser. B Methodol. 57 (1995) 289–300. 10.1111/j.2517-6161.1995.tb02031.x.

[36] K. Wijesooriya, S.A. Jadaan, K.L. Perera, T. Kaur, M. Ziemann, Urgent need for consistent standards in functional enrichment analysis, PLOS Comput. Biol. 18 (2022) e1009935. 10.1371/journal.pcbi.1009935.

[37] P.B. Petrov, J.M. Considine, V. Izzi, A. Naba, Matrisome AnalyzeR – a suite of tools to annotate and quantify ECM molecules in big datasets across organisms, J. Cell Sci. 136 (2023) jcs261255. 10.1242/jcs.261255.

[38] Y. Perez-Riverol, C. Bandla, D.J. Kundu, S. Kamatchinathan, J. Bai, S. Hewapathirana, N.S. John, A. Prakash, M. Walzer, S. Wang, J.A. Vizcaíno, The PRIDE database at 20 years: 2025 update, Nucleic Acids Res. 53 (2025) D543– D553. 10.1093/nar/gkae1011.

[39] J.M. Redwine, C.F. Evans, Markers of Central Nervous System Glia and Neurons In Vivo During Normal and Pathological Conditions, in: B. Dietzschold, J.A. Richt (Eds.), Prot. Pathol. Immune Responses CNS, Springer, Berlin, Heidelberg, 2002: pp. 119–140. 10.1007/978-3-662-09525-6_6.

[40] R. De Gioia, F. Biella, G. Citterio, F. Rizzo, E. Abati, M. Nizzardo, N. Bresolin, G.P. Comi, S. Corti, Neural Stem Cell Transplantation for Neurodegenerative Diseases, Int. J. Mol. Sci. 21 (2020) 3103. 10.3390/ijms21093103.

[41] I. Ladran, N. Tran, A. Topol, K.J. Brennand, Neural stem and progenitor cells in health and disease, Wiley Interdiscip. Rev. Syst. Biol. Med. 5 (2013) 701–715. 10.1002/wsbm.1239.

[42] D.A. Sufieva, D.E. Korzhevskii, Proliferative Markers and Neural Stem Cells Markers in Tanycytes of the Third Cerebral Ventricle in Rats, Bull. Exp. Biol. Med. 174 (2023) 564–570. 10.1007/s10517-023-05748-8.

[43] F. M, Regulation of secretory vesicle traffic by Rab small GTPases, Cell. Mol. Life Sci. CMLS 65 (2008). 10.1007/s00018-008-8351-4.

[44] Y. Homma, S. Hiragi, M. Fukuda, Rab family of small GTPases: an updated view on their regulation and functions, FEBS J. 288 (2021) 36–55. 10.1111/febs.15453.

[45] H. Stenmark, Rab GTPases as coordinators of vesicle traffic, Nat. Rev. Mol. Cell Biol. 10 (2009) 513–525. 10.1038/nrm2728.

[46] F.R. Kiral, F.E. Kohrs, E.J. Jin, P.R. Hiesinger, Rab GTPases and Membrane Trafficking in Neurodegeneration, Curr. Biol. 28 (2018) R471–R486. 10.1016/j.cub.2018.02.010.

[47] A.M. Craig, M. Jareb, G. Banker, Neuronal polarity, Curr. Opin. Neurobiol. 2 (1992) 602–606. 10.1016/0959-4388(92)90025-G.

[48] H. Park, M. Poo, Neurotrophin regulation of neural circuit development and function, Nat. Rev. Neurosci. 14 (2013) 7–23. 10.1038/nrn3379.

[49] A. Dravid, S. Parittotokkaporn, Z. Aqrawe, S.J. O’Carroll, D. Svirskis, Determining Neurotrophin Gradients in Vitro To Direct Axonal Outgrowth Following Spinal Cord Injury, ACS Chem. Neurosci. 11 (2020) 121–132. 10.1021/acschemneuro.9b00565.

[50] J.R. Murillo, L. Goto-Silva, A. Sánchez, F.C.S. Nogueira, G.B. Domont, M. Junqueira, Quantitative proteomic analysis identifies proteins and pathways related to neuronal development in differentiated SH-SY5Y neuroblastoma cells, EuPA Open Proteomics 16 (2017) 1–11. 10.1016/j.euprot.2017.06.001.

[51] L. Goto-Silva, M. Martins, J.R. Murillo, L.R.Q. Souza, G. Vitória, J.T. Oliveira, J.M. Nascimento, E.C. Loiola, F.C.S. Nogueira, G.B. Domont, M.Z.P. Guimarães, F. Tovar-Moll, S.K. Rehen, M. Junqueira, Quantitative profiling of axonal guidance proteins during the differentiation of human neurospheres, Biochim. Biophys. Acta BBA - Proteins Proteomics 1869 (2021) 140656. 10.1016/j.bbapap.2021.140656.

[52] N. Marotta, S. Kim, D. Krainc, Organoid and Pluripotent Stem Cells in Parkinson’s Disease Modeling: An Expert View on their Value to Drug Discovery, Expert Opin. Drug Discov. 15 (2020) 427–441. 10.1080/17460441.2020.1703671.

[53] J. Cerneckis, G. Bu, Y. Shi, Pushing the boundaries of brain organoids to study Alzheimer’s disease, Trends Mol. Med. 29 (2023) 659–672. 10.1016/j.molmed.2023.05.007.

[54] E.C. Ko, S. Spitz, F.M. Pramotton, O.M. Barr, C. Xu, G. Pavlou, S. Zhang, A. Tsai, A. Maaser-Hecker, M. Jorfi, S.H. Choi, R.E. Tanzi, R.D. Kamm, Accelerating the in vitro emulation of Alzheimer’s disease-associated phenotypes using a novel 3D blood-brain barrier neurosphere co-culture model, Front. Bioeng. Biotechnol. 11 (2023). 10.3389/fbioe.2023.1251195.

[55] X. Shao, C.D. Gomez, N. Kapoor, J.M. Considine, C. Grams, Y. (Tom) Gao, A. Naba, MatrisomeDB 2.0: 2023 updates to the ECM-protein knowledge database, Nucleic Acids Res. 51 (2023) D1519–D1530. 10.1093/nar/gkac1009.

[56] M. Pavel, M. Renna, S.J. Park, F.M. Menzies, T. Ricketts, J. Füllgrabe, A. Ashkenazi, R.A. Frake, A.C. Lombarte, C.F. Bento, K. Franze, D.C. Rubinsztein, Contact inhibition controls cell survival and proliferation via YAP/TAZ-autophagy axis, Nat. Commun. 9 (2018) 2961. 10.1038/s41467-018-05388-x.

[57] R. Villanueva, Advances in the knowledge and therapeutics of schizophrenia, major depression disorder, and bipolar disorder from human brain organoid research, Front. Psychiatry 14 (2023) 1178494. 10.3389/fpsyt.2023.1178494.

[58] L. Goto-Silva, M. Junqueira, Single-cell proteomics: A treasure trove in neurobiology, Biochim. Biophys. Acta BBA - Proteins Proteomics 1869 (2021) 140658. 10.1016/j.bbapap.2021.140658.

