## Supplementary material for "A fast TMT-based proteomic workflow reveals neural enrichment in neurospheres of hiPSC-derived neural stem cells"

**Suplementar material**

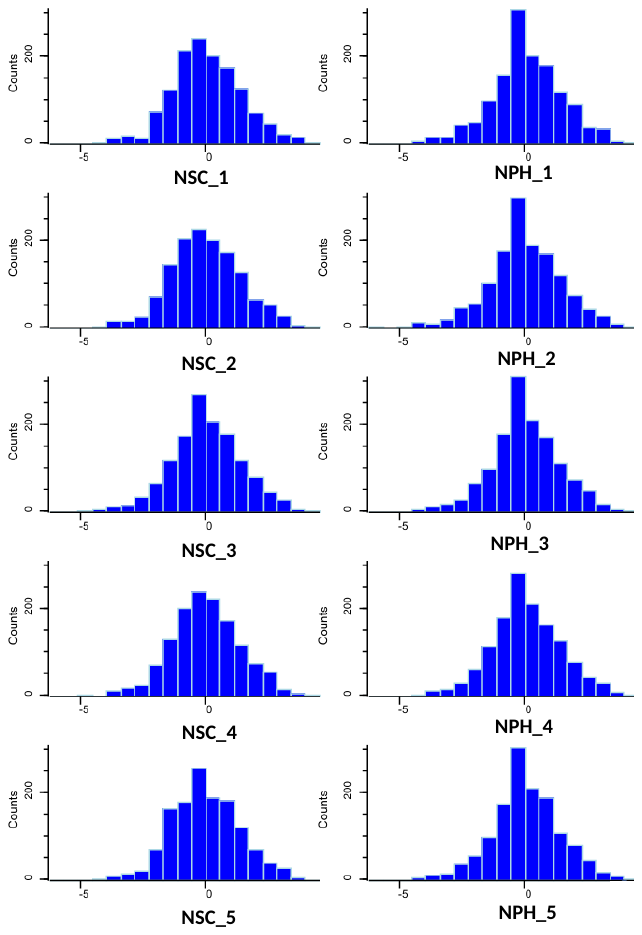

**Suplementar fig 1.** Histograms of normalized abundance values of each sample. Each sample was labeled with an exclusive TMT reagent (which means an exclusive reporter ion signal, or “channel”, represents each sample). The relative abundance values of each reporter ion were normalized in Perseus software by transforming in log_2_ and subtracting by the median. No normalization issue is found in any sample.

**Links to the original free images from Scidraw and Servier Medical Art (used in our workflow and graphical abstract):**

1 – <https://beta.scidraw.io/drawing/444>

2 – <https://beta.scidraw.io/drawing/532> (we altered the colors of this one)

3 – <https://beta.scidraw.io/drawing/300>

4 – <https://beta.scidraw.io/drawing/736>

5 – <https://smart.servier.com/smart_image/stem-cell/>

6 – <https://smart.servier.com/smart_image/normal-cell-cancer/>

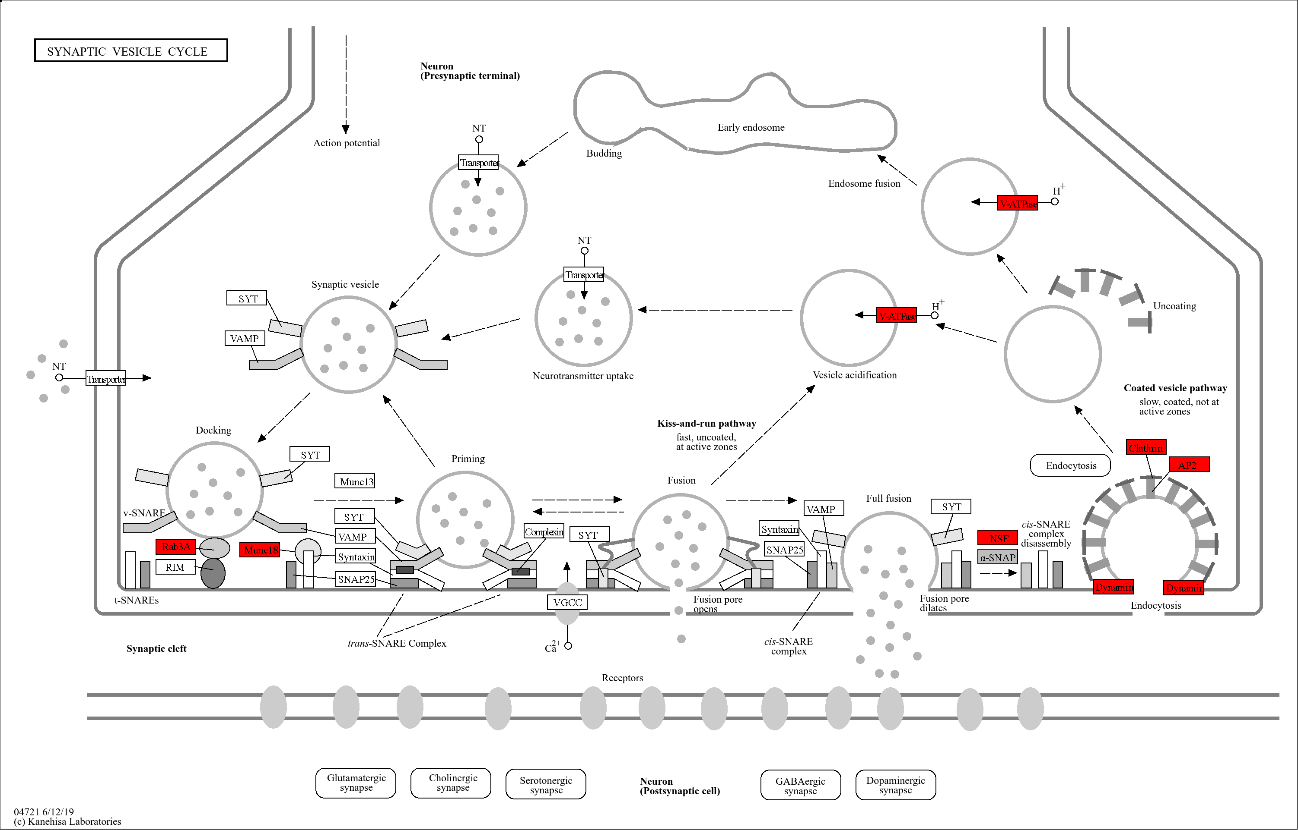

**Suplementar fig 2.** Coverage of the Kegg pathway “Synaptic Vesicle Cicle”. The proteins in red had increased abundance in NPHs, the ones in white were not identified and the protein α-SNAP (in grey) was not differential. The goal of this analysis is to assess how much of the proteins from a relevant pathway like this one can we identify with our method. We can see that there was a considerable part of the pathway we could not cover. This figure was made with the “Color” function provided by Kegg web tools.
